# The Brain/MINDS 3D digital marmoset brain atlas

**DOI:** 10.1101/228676

**Authors:** Alexander Woodward, Tsutomu Hashikawa, Masahide Maeda, Takaaki Kaneko, Keigo Hikishima, Atsushi Iriki, Hideyuki Okano, Yoko Yamaguchi

## Abstract

We present a new 3D digital brain atlas of the non-human primate, common marmoset monkey (*Callithrix jacchus*), with MRI and coregistered Nissl histology data. To the best of our knowledge this is the first comprehensive digital 3D brain atlas of the common marmoset having normalized multi-modal data, cortical and sub-cortical segmentation, and in a common file format (NIfTI). The atlas can be registered to new data, is useful for connectomics, functional studies, simulation and as a reference.

The atlas was based on previously published work but we provide several critical improvements to make this release valuable for researchers. Nissl histology images were processed to remove illumination and shape artifacts and then normalized to the MRI data. Brain region segmentation is provided for both hemispheres. The data is in the NIfTI format making it easy to integrate into neuroscience pipelines, whereas the previous atlas was in an inaccessible file format. We also provide cortical, mid-cortical and white matter boundary segmentations useful for visualization and analysis.

## Background & Summary

A brain atlas provides a reference for the anatomical brain regions of a particular species of study. Accordingly, neuroscientific progress is reflected in the refinement and evolution of brain atlases published throughout the ages. Nowadays, the emergence of digital brain atlasing provides a number of benefits over the traditional book form; we can generate arbitrary cutting planes through the data, no longer limited to a presentation in 2D along some predetermined anatomical plane, and a digital atlas is amenable to computational processing and transformation, forming a critical component of quantitative analysis pipelines on 3D brain data.

In this data report we present a new 3D digital atlas of the non-human primate, common marmoset (*Callithrix jacchus*), a new world monkey that has become an increasingly popular research species within biology and neuroscience as a brain disease model Okano et al.^10^. (Throughout this article ‘marmoset’ shall refer to the common marmoset.) The atlas was constructed as part of Japan’s Brain/MINDS project which aims to map the marmoset brain connectome at micro, meso, and macro-scales, from both structural and functional perspectives - see Okano et al.^10^ for an overview. The brain atlas of this report was developed due to the absence of a comprehensive 3D digital atlas in a format that can be easily used with a wide range of tools for neuroinformatics data integration and analysis. The atlas is currently being applied to connectomics studies (DWI and tracer injection), functional studies (fMRI), as a common space for data integration, and for brain simulation. Summarily, the 3D digital atlas plays a central role in creating a comprehensive knowledge space by integrating the results of a number of different experiments.

As an object of neuroscientific study the marmoset has a number of attractive features. Summarizing Hardman and Ashwell^4^: it is a primate and compared to rodents, is neurologically and genetically closer to humans, and has a more complicated social behavior. Marmosets breed more rapidly than larger primates and have shorter lifespans; helpful for long-term studies of neurodegenerative diseases such as Alzheimer’s and Parkinson’s. And their smaller size compared to primates such as macaques make them easier to handle.

The marmoset brain is on average around 30mm in length and 20mm in width with a weight around 8g (Solomon and Rosa^14^). The cortical surface is generally smooth except for regions of the visual cortex located in the banks of the calcarine sulcus^14^ and the lateral fissure (whose depth and extent varies reasonably between individuals). Furthermore, marmoset and human brains share a particularly subdivided frontal lobe, expanded temporal lobe, and also striate cortex dependent visual pathways^12^.

A number of marmoset brain atlases have been published, but the majority of them are in article or book form instead of digital. Also, none of them have precisely co-registered Nissl and MRI data in a digital format with both cortical and subcortical structures.

In chronological order: Saavedra and Mazzuchelli^13^ (a hard to find article), Stephan et al.^15^ (an out of print book), Palazzi and Bordier^11^, Yuasa et al.^18^ (book and also available in an online version), Hardman and Ashwell^4^ (book), Paxinos et al.^12^ (a book that is commonly used as the standard reference). Digital versions of the cortical regions (for one hemisphere) of the Paxinos et al.^12^ atlas can be found at a number of sites online, e.g. Majka and Kowalski^8^, or from *The Scalable Brain Atlas* website (Bakker et al. ^2^). The original Nissl histology images are also available online at Tokuno et al.^16^. The cortical parcellation and subsampled Nissl data can be downloaded as supplementary material to Majka et al.^9^.

Finally, a digital atlas was described in Hashikawa et al.^6^ and is downloadable from http://brainatlas.brain.riken.jp/marmoset/modules/xoonips/listitem.php?index_id=66. This data can be viewed using the proprietary *SgEye*-*Viewer* software. The atlas was constructed using a cytoarchitectonic approach through Nissl histological staining and region delineation of 86 brain slices. The aforementioned publication and digital data support a forthcoming book based on the same data (Hashikawa et al.^5^). The book version includes axial, coronal and parasagittal slices of the Nissl and MRI showing region overlays. Regions are delineated as in the digital version along with the additional labeling of small regions (without boundary specification), specified in both the Nissl and MRI. (This was due to the technical difficulty in specifying such small regions in the digital model.) In both the digital and book forms the regions were delineated for only the left hemisphere. Currently, the Human Brain Project (HBP) also hosts the 86 axial Nissl slices (no MRI data) with annotation in the left hemisphere and can be viewed in their web-based atlas viewer: https://nip.humanbrainproject.eu/atlas/#/?dataset=67d4b403-77b3-4955-9926-07c59c28a66a. The digital version of this atlas was created and saved in a proprietary data format, only displayable from the SgEye-Viewer software. One of the main difficulties with this atlas release is that the custom data format makes it hard to use in neuroinformatics pipelines.

Our work relates to this atlas and book in that the same original Nissl and MRI data was used as the basis for our new atlas. We went back to the original dataset and used state-of-the-art computational processing to create a more accessible and multi-modal brain atlas with higher fidelity in 3D. We improve on the previous Hashikawa et al.^6^ atlas by removing inter-slice illumination and shape artifacts, and normalizing the reconstructed Nissl volume to the MRI data shape. Furthermore, originally only 86 axial Nissl histological images were annotated, even though a larger number of brain sections were taken. So we used double the number of Nissl images (172) to create our Nissl volume and removed inter-slice deformation between them. Resulting in a smoother and more detailed atlas. The Nissl and parcellation volumes were then normalized to the MRI shape using 3D image registration techniques.

The brain region segmentation was mirrored into the right hemisphere in order to provide a complete parcellation over the entire brain, something very useful for connectomics. Summarily, this data descriptor focuses on the new computational work carried out in order to process this data.

The normalization of the histology and parcellation data to the MRI shape was chosen as MRI data for each individual brain can be taken non-invasively and can serve as the common modality for data registration and integration in large-scale brain mapping projects.

The second important aspect of this data release is that the atlas is provided in the common NIfTI format, with separate volumes for the MRI, Nissl and region segmentation (parcellation), whereas the previous work was in a proprietary format that could only be loaded into custom software. Therefore, this atlas can be easily loaded and integrated into neuroscience tools and pipelines, or directly manipulated via programming languages such as Python or Matlab.

To help introduce potential users to the atlas a scene file for the freely available and open-source 3D Slicer software (https://www.slicer.org) is also included. Here the user can interact with the atlas in 3D and view or overlay the region annotations, MRI and Nissl data interactively. The data will appear as in Fig. 1 when loaded.

**Figure 1.**
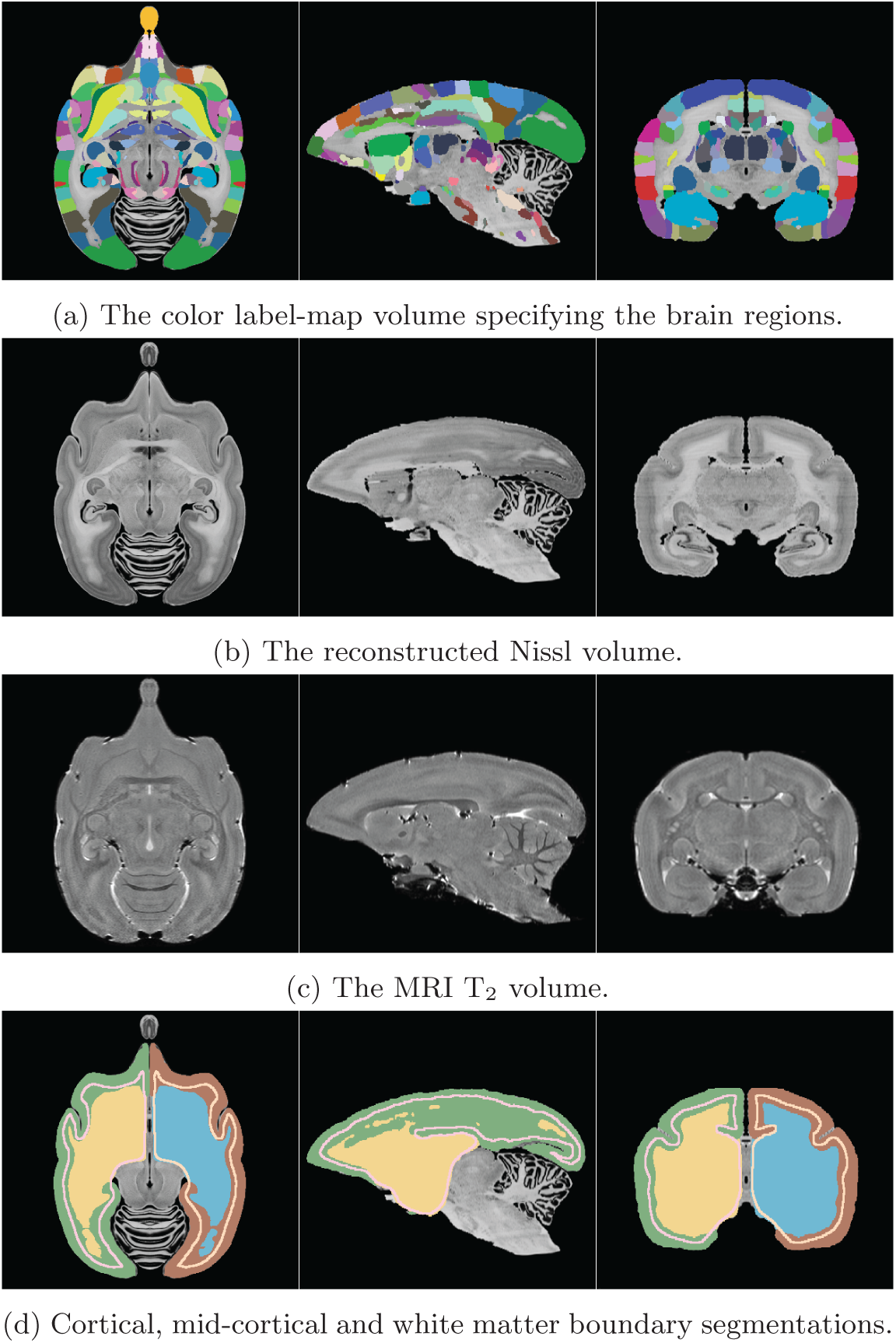
The new atlas – axial, sagittal and coronal views of the (a) region segmentation, (b) Nissl, (c) MRI and (d) gray/mid/white matter boundary segmentation volumes. Note the similarity in shape between the Nissl and MRI due to the Nissl being normalized to the MRI shape. The colors of the region segmentation are symmetric between hemispheres but each region has a unique ID, described in the provided lookup table. The gray, mid-cortical and white matter boundary segmentation is displayed using the default color lookup settings of 3D Slicer, showing unique colors for each hemisphere.

To summarize, Fig. 2 gives an overview of the data generation workflow for the atlas, and a precise description is given in the *Methods* section.

**Figure 2.**
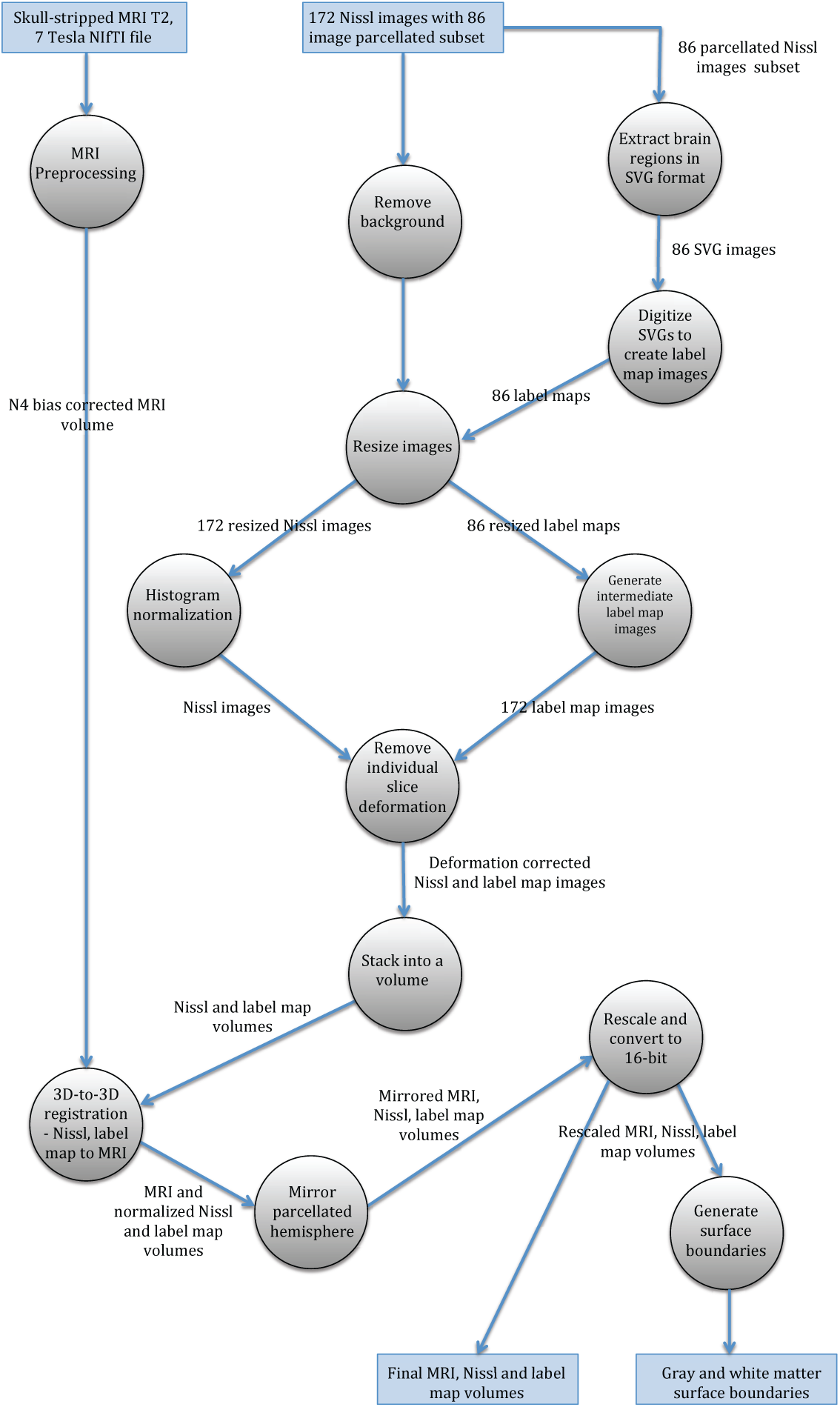
Overview of data generation workflow by using a dataflow diagram. The circles represent processes and their description is given in the Methods section.

The dataset described in this report is available at the Brain/MINDS dataportal (Data Citation 1). The data can be downloaded as individual files or together as one.zip file.

## Methods

In this section we first give an overview of how the brain image data was acquired. We then give an in-depth description of the data processing that was used to create this data release.

## Data acquisition

An ex-vivo MRI *T*_2_ weighted image of an adult marmoset (detailed in Table 1) was firstly acquired ( 7-T PharmaScan 70/16 MRI scanner (Bruker biospec MRI GmbH; Ettlingen, Germany)). Based on brain image and histology analysis there were no abnormalities detected. For behavior and physical condition, also no abnormality was detected.

**Table 1.** Marmoset specimen details.

|  |  |
| --- | --- |
| Species | Common Marmoset (*Callithrix jacchus*) |
| Sex | Female |
| Age | 2.3 years old |
| Weight | 310 g |

Following perfusion the skull with brain was mounted on a stereotaxic apparatus with carbon shafts inserted into the brain to create reference marks visible in the MRI image. The brain was then embedded in gelatin, sliced along the axial plane at 50 μm intervals, and these slices mounted onto slides. The slices were stained with thionine and photomicrographs were acquired using an Olympus VS-100 virtual slide scanner.

## Brain region segmentation

The brain region segmentation was performed in a previous study to this work and is presented in Hashikawa et al.^6^ and available at http://brainatlas.brain.riken.jp/marmoset/modules/xoonips/listitem.php?index_id=66. The histology images, stored as TIF image files (.tif extension), were first manually aligned and stacked into an initial volume. Then, virtual coronal sections were generated from this data and the Paxinos et al.^12^ atlas was used as a reference to create a first coronal plane cortical delineation. This delineation was then transferred back to the original data (axially cut slices) and confirmed through further histological examination. Sub-cortical regions were annotated in a corresponding manner ^6^.

Delineation of brain regions was done using Photoshop and each region boundary was stored as a Photoshop *path* and given a name based on abbreviations consistent with the Paxinos et al.^12^ atlas. All *paths* for a slice were stored inside the corresponding TIF file. External to the TIF files a text file was used to describe the region name, abbreviation and a color. This process was done for a sub-set of 86 slices – every fourth slice of the original data.

Aligning and stacking this set of 2D images gives a 3D volume with segmented (parcellated) regions. The TIF images from this manual procedure were used as input to our data processing.

## Data processing summary

A number of improvements can be made to the quality of the data described in the previous section. Firstly there is illumination variation between slices. Secondly, there is slice deformation due to the slide mounting and staining process which can be improved upon by computationally enforcing inter-slice consistency. We can also integrate more slices into the atlas and upsample the brain region parcellation to this resolution to give a higher fidelity image along arbitrary cutting planes.

Furthermore, in order to create a multi-modal atlas, the 3D shape of the Nissl data should be normalized to a chosen reference modality. As mentioned before, the ex-vivo MRI T_2_ data of the same animal was used as the reference modality.

The atlas in this data report was created by going back to the source TIF files and applying the aforementioned improvements by using a more advanced processing workflow. A set of manually aligned Nissl stained images were stacked (172 slices) and high-frequency distortions between histology slices were removed using an approach based on Gauss-Seidel iteration^3^ and using the ANTs normalization tools (version 2.1.0)^1^. Every second TIF image had brain region segmentation information (i.e. 86 slices); this was interpolated into a 172 slice resolution and turned into digital format in the form of a segmentation volume where each voxel has an ID corresponding to a brain region.

Then 3D-to-3D registration between the Nissl and MRI was performed by combining landmark based registration in the 3D Slicer software and automatic registration using ANTs. Landmarks were included into the registration workflow to help the optimization algorithm reach a strong solution of the objective function for image matching. Then, the calculated transformations were applied to the region segmentation volume to generate the final atlas parcellation. For all relevant steps we used ANTs’ SyN algorithm for nonlinear registration, considered to be one of the top performers for biomedical image registration^7^.

In summary, the input data for the creation of this data release was as follows:

1. 172 TIF images of axially sliced Nissl stained sections.
  a. 86 of these images (every second image) with region segmentation in Photoshop *path* format.
  b. Resolution: 8000 × 5420 pixels.
2. 210 TIF images of MRI data with non-brain data removed.
  a. Resolution: 468 × 360 pixels.
3. A NIfTI version of the MRI data with non-brain data included.
  a. Resolution: 226 × 512 × 186 pixels. With a voxel size of 0.150 mm × 0. 100 mm × 0.150 mm.
4. A text file listing brain regions, giving parent/child structure, name, abbreviation and color.

We now give a precise description of each of the processing steps as overviewed and as shown in Fig. 2.

## Data processing step-by-step description

### MRI data processing

The skull-stripped MRI data that was received as a set of TIF files was converted to a single NIfTI 3D volume file (.nii). The correction physical proportions for this data were found by matching with the reference non-skull-stripped MRI NIfTI data that was also provided. The N4 bias correction algorithm^17^ in 3D slicer was used to correct intensity inhomogeneities in the MRI T_2_ data (such artifacts can result due to the nature of the MRI imaging system). We also padded the MRI volume above and below (along the S axis) as registration algorithms have trouble treating data at the boundaries of an image volume. This becomes important when the atlas is to be registered to new data.

### Atlas coordinate system

The MRI was used as the reference space for the brain atlas, therefore the brain orientation and reference coordinate system had to be defined. The RAS coordinate system (Talairach-like), common to the 3D Slicer software (see http://wiki.slicer.org/slicerWiki/index.php/Coordinate_systems for a description), was chosen: X-axis from Left to (R)ight, Y from Posterior to (A)nterior, Z from Inferior to (S)uperior.

The brain was oriented to align with the stereotaxic coordinate system defined in Paxinos et al.^12^ by reference to the digital dataset accompanying Majka et al.^9^. This makes it very easy to take comparable measurements in 3D between the two atlases. Here the left-right zero plane coincides with the midsagittal plane, the axial (horizontal) zero plane is the plane ‘passing through the lower margin of the orbit and the center of the external audiotory meatus’^12^. Figures 3 and 4 describe the atlas coordinate system.

**Figure 3.**
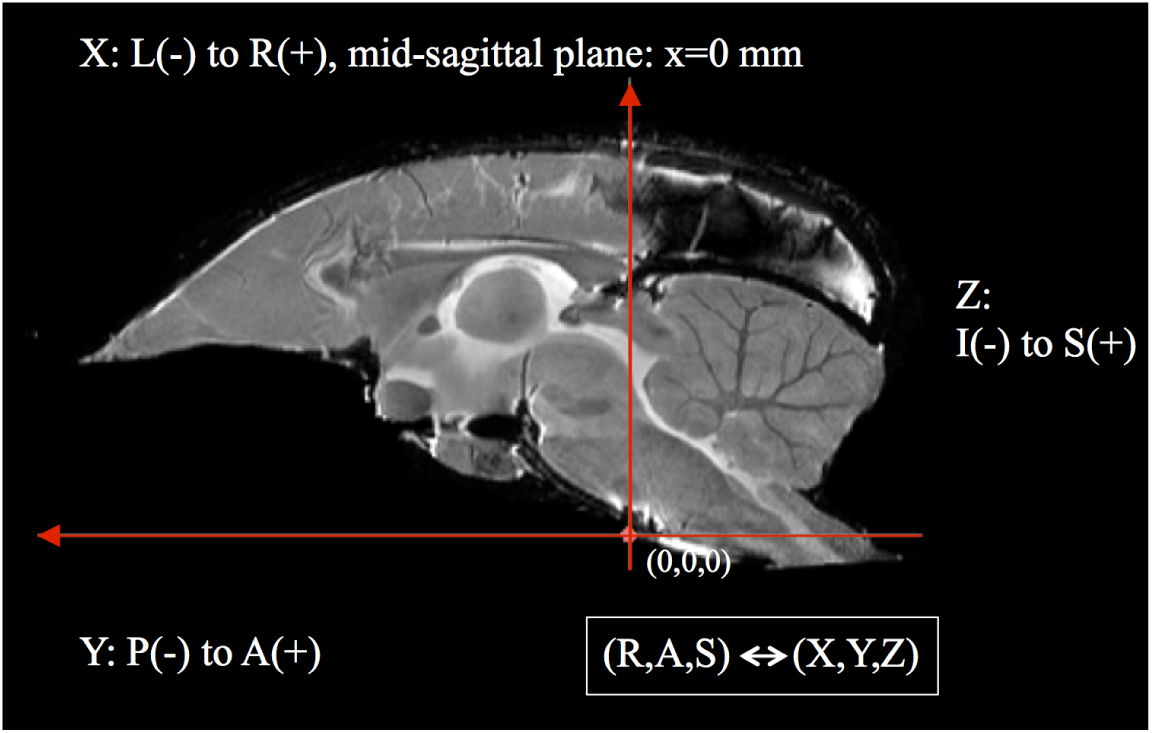
A diagramatic mid-sagittal view of the atlas showing the ac-pc alignment and RAS coordinate system mapping with XYZ.

**Figure 4.**
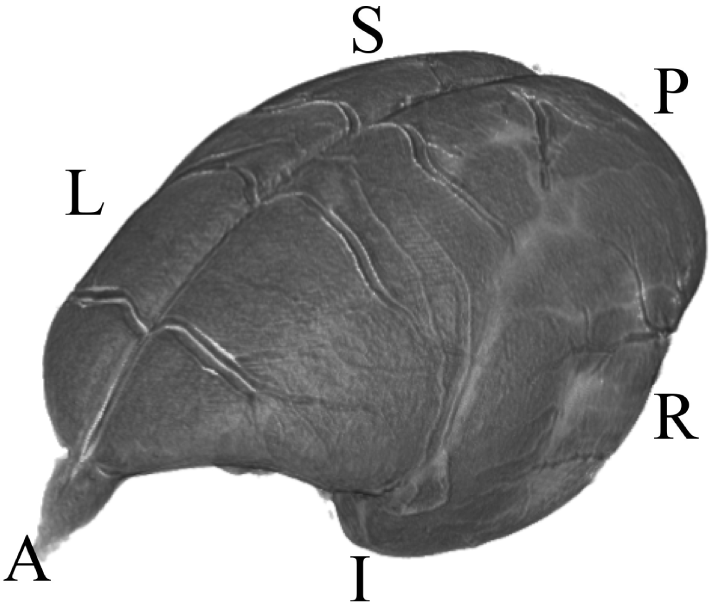
A 3D view depicting the RAS coordinate system and the orientation of the brain within it. The letters denote the orientation as follows: (A)nterior, (P)osterior, (S)uperior, (I)nferior, (R)ight, (L)eft.

### Extract brain region information from Nissl histology images

The histology TIF images were received as two sets: one with cortical Photoshop *paths* defining the boundaries of anatomical regions, and the other with sub-cortical ones. We processed these images by exporting all *paths* into the SVG format (Scalable Vector Graphics, defined as an XML text file) along with the *path* name (set to an anatomical region’s abbreviation). Then all of the SVGs for a particular histology image were consolidated into one concatenated SVG file. During this process SVGs were cross-referenced with the list of regions given separately in a text file, checking for any unused or extraneous regions that are not part of the final list. Each region was assigned a unique ID and corresponding unique temporary color to each region (due to the final assigned colors being non-unique). This information was saved in a text file in preparation for digitizing the SVG files into digital 2D region segmentation images.

### Digitize 2D brain regions and create 2D segmentation images

The 86 consolidated SVG files were loaded into Adobe Illustrator and then converted and saved as RGB color raster images. Then, each pixel value was changed from the temporary unique RGB color to the reference region ID. To support a wide number of ID values the images were saved as single channel 16-bit TIF files.

### Nissl background removal

The original histology images had a white background. For each image this region was removed and set to black by using a mask based on the Adobe Photoshop *path* representing the whole brain region.

### Resize Nissl and label map images

Both the Nissl histology images and corresponding region segmentation images were resized then reoriented by 90 degrees to align with RAS orientation in anticipation of stacking into a volume, giving an image size of 678 × 1000 pixels. This size was chosen as a compromise between image resolution and computational tractability in 3D-to-3D nonlinear registration.

### Resample 2D region segmentation images

At this stage of processing there were 172 Nissl histology images and region segmentation for only 86 of these images (every second image). We generated 172 region segmentation images by stacking the 86 images into a tentative 3D volume with the correct voxel spacing, upsampling the volume to a reference space with double the resolution in the S-axis (172 slices), and then extracting 172 axial images from this volume. The ANTs software and its *MultiLabel* interpolation method was used to do this. (*MultiLabel* interpolation was used for all transformation steps applied to region segmentation data in order to preserve integer labeling.)

### Nissl image histogram normalization

Non-uniform illumination between histological image slices can occur due to strobe lighting or through the use of a camera’s auto-white balance control. Additionally, the nature of the Nissl staining procedure means that the intensity of the stain can vary between slices. If the image set is stacked into a volume and the volume is viewed coplanar to the cutting plane this variation will show up as intensity banding across the brain image.

We treated this variation by using a histogram normalization procedure. For each image in the stack we normalized its histogram to its nearest neighbors (handling the first and last images as special cases) and then averaged the result to get a new adjusted image. This procedure was done for three iterations to successfully ameliorate the illumination variation.

### Removing individual slice deformation from histology and region segmentation images

The set of histological slices had independent local deformations caused by the cutting, slide mounting, and staining procedure. Removing this deformation was an important step for improving the atlas from what was previously available, constructing a volume where fine structures appear consistent as they continue across adjacent slices.

Deformation removal was achieved through an iterative Gauss-Seidel approach, described in Gaffling et al.^3^. Using this approach we can iteratively attenuate high-frequency noise (manifest in the small inconsistencies in shape between slices) while preserving low-frequency information (the brain shape). The basic idea was that for each slice, ANTs was used to perform non-linear registration between it and its neighbors. These transformations were then used to update the slice’s shape. This process was performed for four iterations, generating a set of 172 transformation files. The improvement in fine detail can be seen in Fig. 5.

**Figure 5.**
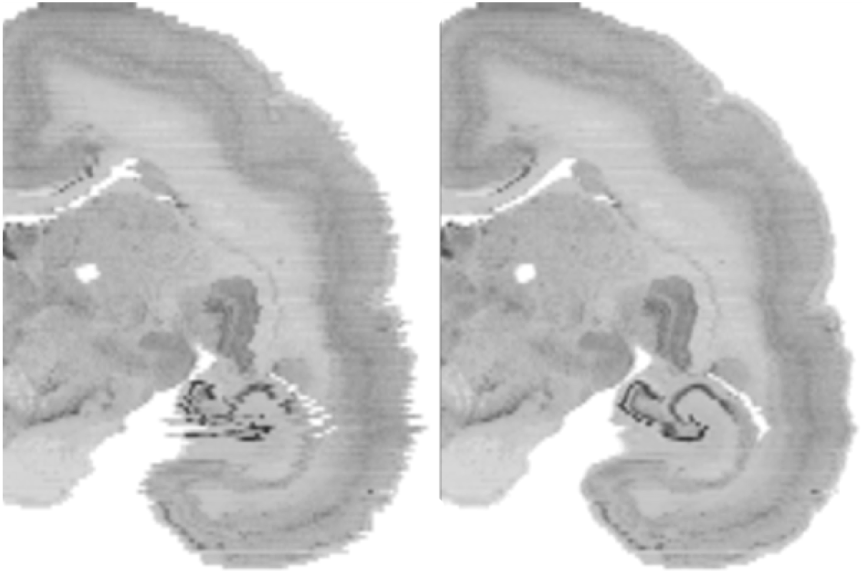
A coronal view of a slice of the brain showing before (left) and after (right) inter-slice distortion removal for the Nissl data. Note the clear improvement in the hippocampal region.

These transformation files were also applied to the corresponding region segmentation images to update their shape. Both 2D Nissl and region segmentation images were then converted to NIfTI (.nii) 3D image volumes and oriented to the RAS standard in order to be used as input for the next step. For both datasets, blank space was added as extra slices above and below (along the S-axis) to give padding to the volume.

### 3D-to-3D registration to normalize the Nissl and region segmentation volume shapes

#### Part 1: Landmark matching in 3D slicer

A consistent Nissl volume was prepared out of the initial data that can now be normalized to the shape of the MRI T_2_ data. Since precise inter-modal image registration is difficult, we took a semi-automated, iterative approach to complete this task by using both landmark and automated image registration. For each step the MRI volume was set as the *fixed* image and the Nissl as the moving image.

Firstly, ANTs linear registration (up to an Affine transformation) was used to initialize the alignment of the Nissl volume to the MRI volume. Next manual landmark based registration was used. This step was necessary as it was found that automatic registration on its own using SyN faced problems in reaching a strong global minima in the image registration. This was due to large differences in contrast mechanism between the MRI and Nissl data, and also due to the gradient descent nature of the algorithm, making it sensitive to initial conditions.

The two volumes were loaded into the 3D Slicer software and its *Landmark Registration* module was used to perform interactive nonlinear registration. Corresponding landmarks were manually set for each dataset and thin-plate spline deformation updated the Nissl volume shape. This interactive approach allowed for visual inspection and validation of the shape deformation. This landmark-based registration was refined until a close as possible match between the MRI and Nissl was achieved. Then the normalized Nissl volume was exported from 3D slicer.

3D Slicer exports the updated volume but does not export any usable transformation file that can be included into the ANTs workflow for updating the region segmentation volume. Therefore we used ANTs SyN to calculate the nonlinear transformation between the Nissl volume before and after landmark registration. I.e. the fixed image was the Nissl volume after landmark registration and the moving image was the volume before. Doing this allowed us to generate a transformation warp field that could be applied to the region segmentation volume.

#### Part 2: Refine the registration with ANTs

The normalization of the Nissl to MRI was refined using the purely automated nonlinear SyN registration algorithm. Since the algorithm works directly on the image data we could refine the matching of structural boundaries in a manner more comprehensive than specifying landmarks alone. Then the calculated transformation warp field was concatenated with that from the previous step.

#### Part 3: Correct the occipital lobe

One of the limitations of the available registration tools is the inherent regularization in the generated transformation fields. In reality there can exist discontinuities in the best map between two imaging modalities of the same brain. In this dataset the Nissl image volume has a gap between the occipital lobe and the cerebellum, whereas in the MRI data this space was closed with the two regions abutting one another. Neither the 3D Slicer nor ANTs SyN algorithms can effectively deal with this registration situation so we used a masking procedure to register the occipital lobe separately and then combined this result with the previously calculated whole brain transformation field.

Specifically, the occipital lobe in both the MRI and Nissl was delineated (excluding the adjacent white matter). These binary masks were then nonlinearly registered one to the other using SyN, with the MRI derived mask as fixed data and the Nissl derived mask as moving data. The calculated transform field was then applied to the Nissl volume (after transformation from the previous step).

In order to combine the transform fields a mask covering the occipital lobe (including the adjacent white matter) in the newly warped Nissl space was created. This mask was then used to replace the transformation warp field data in the global transformation with the correct occipital lobe specific transformation data. A weighted blending between the global and occipital lobe specific transforms was used in the region where the occipital lobe met with the rest of the cerebral cortex. Finally, this newly constructed transformation was applied to the Nissl data and region segmentation volume, giving the final normalization of the data to the MRI shape. The base resolution of the data was standardized for all modalities as a compromise between data resolution and computational friendliness.

#### Mirror hemispheres

The original brain data was annotated for one hemisphere only so this was mirrored in order to create a complete atlas. Since the MRI data was already axially aligned (as described earlier in this section) it was trivial to cut and mirror the data across the mid-sagittal plane. This was repeated for the Nissl and region segmentation volumes. Additional processing was applied to the region segmentation where the value of 10000 was added to every brain region ID in the left hemisphere in order to differentiate between the two hemispheres. Further information on ID assignment is given in the data records section.

#### Rescale and convert voxel data to 16-bit

The calculation of all transformations was done with floating point data but this was unnecessary for the number of pixel intensit levels in the original data. We therefore converted all of the data to 16-bit single channel volumes. Additionally, for visual appearance, we rescaled the MRI and Nissl data to fit the dynamic range of a 16-bit single channel image. The region segmentation values, specifying the integer IDs, were left unchanged.

#### Color look-up table

A color look-up table (saved as a text file) was created to assign the region name and color to each unique ID within the region segmentation volume. The format of one line of the is: *ID name R G B A*, where *ID* is an integer, *name* a string, and *RGBA* are 0-255 integers (https://www.slicer.org/wiki/Documentation/4.0/Modules/Colors. This simple file can be loaded directly into 3D Slicer. For display purposes the colors of the region segmentation are symmetric between hemispheres but each region has a unique ID to distinguish between left and right, described in the lookup table.

#### Gray matter, white matter and mid-cortical boundary segmentation creation

This dataset includes segmentations specifying the csf-gray matter boundary, the gray-white matter boundary, and a mid-cortical boundary for both hemispheres. This data is provided as NIfTI voxel data and can be easily turned into a polygonal surface format for use in various types of analysis and visualization. For example, the white matter segmentation could be used as a seed region for diffusion MRI and the mid-cortical surface can be extracted from the volume and used for projecting and visualizing neocortical myelin density, or for functional statistics maps for fMRI experiments.

The csf-gray boundary and gray-white boundary were created by manual parcellation. These were used as input to the freely available FreeSurfer software tools (https://surfer.nmr.mgh.harvard.edu/) in order to automatically identify a mid-cortical layer (created by FreeSurfer’s *mris*_*make*_*surfaces*). This was then exported and rasterized into a digital volume, then manually corrected for errors in 3D Slicer.

## Code availability

Due to the complexity and heavy manual aspect of the data creation, the code used to carry out particular steps is not provided. However, we believe that the precise description of the data generation given in the methods section of this article should be useful for anyone wanting to create a similar atlas and the authors can be contacted for any further questions regarding atlas construction.

## Data Records

This work contains a single data record containing a set of volume files in NIfTI format for the MRI, Nissl, and segmentations. All NIfTI files were saved with compression (.nii.gz extension) to reduce the file size. Additionally a color look up table to match voxel IDs with brain regions and a color is given. The description of their generation was given in the previous *Methods* section. Lastly, a 3D slicer scene file is given so that users can quickly and easily load and test out the atlas in the 3D slicer software. Descriptions of the files and their formats are given in Tables 2 and 3.

**Table 2.** Data file details for this data descriptor.

| Filename | File type | Details |
| --- | --- | --- |
| bma-1-mri.nii.gz | NIfTI | MRI T<sub>2</sub> volume data |
| bma-1-nissl.nii.gz | NIfTI | Nissl volume data |
| bma-1-region_seg.nii.gz | NIfTI | Region segmentation volume data |
| bma-1-graywhite_seg.nii.gz | NIfTI | Gray and white matter boundaries |
| bma-1-mid_seg.nii.gz | NIfTI | Mid-cortical boundary |
| bma-1-clut.ctbl | Text | Color lookup table for region segmentation |
| bma-1-slicerscene.mrml | MRML file | A 3D Slicer scene file for the dataset |
| bma-1-all.zip | ZIP file | A zip file containing all of the above files |

**Table 3.** NIfTI volume file details.

|  |  |
| --- | --- |
| Bit-depth | 16 bits per voxel |
| Volume size | $678 \times 1000 \times 210$ voxels |
| Voxel spacing | $0.040 \text{ mm} \times 0.040 \text{ mm} \times 0.112 \text{ mm}$ voxels |
| File size | 284.8 MB uncompressed (1-38.7 MB compressed), (HFS+ file system) |

Overall the atlas region segmentation contains 280 brain region annotations with (114 cortical) per hemisphere, giving a total of 560 brain region annotations in total. However, for completeness, the list of brain regions defined in the color lookup table is 786 regions per hemisphere (for completeness). Subsequently, the region IDs in the right hemisphere range from 1-786 and in the left hemisphere and 10001-10786, with 0 denoting the background. Users should check the IDs in the region segmentation volume and match this with the color lookup table entry to determine the available annotated brain regions.

## Technical Validation

The validity of the brain region annotation found in this dataset is support by the fact that its raw data was processed (in a different way) and already published in Hashikawa et al.^6^, which also follows the region descriptions and naming conventions in Paxinos et al.^12^.

Furthermore, the normalization of the Nissl volume shape to the MRI shape was performed using the state-of-the-art in image registration tools, namely ANTs, which has been evaluated in papers such as Klein et al.^7^.

In order to quantitatively assess the registration accuracy of the atlas, a small number of robust brain landmarks were manually located in both the MRI and Nissl datasets and their Euclidean distance compared. The landmarks are given in Table 4.

**Table 4.** Landmark descriptions. The abbreviations are as follows: ac, anterior commissure; pc, posterior commissure; cc, corpus callosum; MB, mammillary body; DLG, dorsal lateral geniculate nucleus; 4V, fastigium of the fourth ventricle.

| Landmark: | Description | Difference |
| --- | --- | --- |
| ac | Mid-sagittal center point at start of ac in coronal view going from anterior to posterior. | 0.132 mm |
| pc | Mid-sagittal center point at start of pc in coronal view going from anterior to posterior. | 0.112 mm |
| cc (start) | Mid-sagittal center point at start of cc in coronal view going from anterior to posterior. | 0.055 mm |
| cc (end) | Mid-sagittal center point at end of cc in coronal view going from anterior to posterior. | 0.162 mm |
| MB (left) | Center of the first appearance of the left MB in coronal view going from anterior to posterior. | 0.116 mm |
| MB (right) | Center of the first appearance of the right MB in coronal view going from anterior to posterior. | 0.133 mm |
| DLG (left) | The point at the first appearance of the left DLG in coronal view, moving from anterior to posterior. | 0.068 mm |
| DLG (right) | The point at the first appearance of the right DLG in coronal view, moving from anterior to posterior. | 0.160 mm |
| 4V (fastigium) | Mid-sagittal point of the fastigium of the fourth ventricle, identified in the sagittal plane. | 0.050 mm |

The nine landmarks were chosen in anatomical regions (abbreviations are based on the Paxinos et al.^12^ nomenclature) as follows: ac, anterior commissure; pc, posterior commissure; cc, corpus callosum; MB, mammillary body; DLG, dorsal lateral geniculate nucleus; 4V, fastigium of the fourth ventricle. The positional difference between manually assigned landmarks is shown in Table 4. Given an average error of approximately 0.110 mm we can calculate an approximation of the expected error in voxels to be 0.11/((0.04 + 0.040 + 0.112)/3) ≈ 1.72, or about two voxels. This value is small enough to assume that it arises from error in the manual assignment of landmarks and is small relative to the total number of voxels that the brain occupies.

The specification of the region delineation for this single brain took a number of years to make by a single anatomist at this level of detail (cortical plus subcortical regions). It is possible to do the same thing for multiple subjects to create a population-average atlas, but for practical reasons, precise segmentation of a number of brains at the same level of detail will be a big undertaking, but something for future work. For example, data from a number of modalities can be integrated into the current atlas for a future data-driven approach to brain atlasing. For this, a starting point is needed and our atlas can be used as the base for revising and creating a new version as more and more information is integrated.

## Usage Notes

The NIfTI data format makes this dataset easy to use with common neuroscientific tools such as AFNI, FreeSurfer, or FSL. It also makes it easy for registering the atlas to new marmoset brain data using an image normalization tool such as ANTs. This leads the way to the atlas’ use in fMRI and connectomics. The data can also be loaded programmatically using Matlab, or with Python by using libraries such as SimpleITK.

For getting started we have provided a 3D slicer scene description file (.mrml) that lets a user easily view and interact with the atlas (tested with 3D Slicer version 4.6.2). This is the recommended way to get up and running with the atlas and 3D Slicer is open source and freely available at https://www.slicer.org. Opening the mrml file in 3D Slicer will load the MRI, Nissl, brain region segmentation, color look-up table and grey/mid/white segmentations into the software with the correct visualization settings. The presentation will be similar to that shown in Fig. 1. For basic usage, a user can hover the mouse over any region and see the annotation, move through the data volume along the RAS axes, and adjust which data is being displayed and overlaid.

## Acknowledgements

This research was supported by the Funding Program for World-Leading Innovative R&D on Science and Technology (FIRST Program), JSPS, and the program for Brain Mapping by Integrated Neurotechnologies for Disease Studies (Brain/MINDS) from the Japan Agency for Medical Research and Development, AMED.

The authors would like to thank Akiya Watakabe and Tetsuo Yamamori at RIKEN BSI and Ken Nakae (Kyoto University) for their discussion on landmarks. We thank Noritaka Ichinohe at RIKEN BSI for his advice on landmarks and on marmoset brain anatomy. We thank Reiko Nakatomi at RIKEN BSI for helping create the Nissl stained images.

## Author Contributions

M.M. extracted the brain region boundaries from the TIF files along with providing files for the region and color naming information. T.K. generated the csf/gray, gray/white and mid-cortical segmentations. H.O. and Y.Y. organized and facilitated the development of this brain atlas. A.W. performed all of the data-processing steps described in the text except for the manual editing of the mid-cortical segmentations which were completed collaboratively with T.K. T.H. and A.I created the Nissl stained images and drew the original brain region boundaries in the TIF files. K.H. created the MRI T_2_ data.

## Data Citations

1. Woodward, A., Hashikawa, T., Maeda, M., Kaneko, T., Hikishima, K., Iriki, A., Okano, H., Yamaguchi, Y. *Brain/MINDS Data Portal* https://doi.org/10.24475/bma.2799 (2017).

